# A synthetic microbial loop for modelling heterotroph-phototroph metabolic interactions

**DOI:** 10.1101/379131

**Authors:** Marco Fondi, Francesca Di Patti

## Abstract

Marine ecosystems are characterized by an intricate set of interactions among their representatives. One of the most important occurs through the exchange of dissolved organic matter (DOM) provided by phototrophs and used by heterotrophic bacteria as their main carbon and energy source. This metabolic interaction represents the foundation of the entire ocean food-web.

Here we have assembled a synthetic ecosystem to assist the systems-level investigation of this biological association. This was achieved building an integrated, genome-scale metabolic reconstruction using two model organisms (a diatom *Phaeodactylum tricornutum* and an heterotrophic bacterium, *Pseudoalteromonas haloplanktis*) to explore and predict their metabolic interdependencies. The model was initially analysed using a constraint-based approach (Flux Balance Analysis, FBA) and then turned into a dynamic (dFBA) model to simulate a diatom-bacteria co-culture and to study the effect of changes in growth parameters on such a system. Finally, we developed a simpler dynamic ODEs system that, fed with dFBA results, was able to qualitatively describe this synthetic ecosystem and allowed performing stochastic simulations to assess the effect of noise on the overall balance of this co-culture.

We show that our model recapitulates known metabolic cross-talks of a phototroph-heterotroph system, including mutualism and competition for inorganic ions (i.e. phosphate and sulphate). Further, the dynamic simulation predicts realistic growth rate for both the diatom and the bacterium and a steady state balance between diatom and bacterial cell concentration that matches those determined in experimental co-cultures. This steady state, however, is reached following an oscillatory trend, a behaviour that is typically observed in the presence of metabolic co-dependencies. Finally, we show that, at high diatom/bacteria cell concentration ratio, stochastic fluctuations can lead to the extinction of bacteria from the co-culture, causing the explosion of diatom population. We anticipate that the developed synthetic ecosystem will serve in the future as a basis for the generation of testable hypotheses and as a scaffold for integrating and interpreting-omics data from experimental co-cultures.

## 1. Introduction

The term microbial loop refers to the trophic pathway of the marine food web responsible for the microbial assimilation of dissolved organic matter (DOM) [2, 17]. A key role in such process is possessed by phytoplankton, the pool of free-floating photosynthetic organisms capable of capturing energy from sunlight and thus transforming inorganic matter into organic matter [12]. Indeed, phytoplankton-derived carbon is one of the most important entry points of the microbial loop, being then converted into microbial biomass by heterotrophic bacteria and afterwards transferred up the food web, as they are predated by organisms at higher trophic levels (zooplankton)[12]. Ecologically speaking, the microbial loop has a tremendous importance given that: i) roughly one-half of the available atmospheric oxygen derives from photosynthetic organisms within the marine microbial loop and ii) it represents the foundation of the entire marine food web [33]. At the same time, this biological process is hard to understand in detail, given that many complex interactions at different cellular levels are known to exist among representatives of the different species. This is a typical feature of marine ecosystems that are constructed around networks that encompass many cellular levels (metabolism, signalling, etc) and that connect every species to many others through different interaction types such as mutualism, competition, and parasitism [1]. A well known association occurring within the microbial loop involves diatoms and bacteria [1]. Diatoms are ubiquitous photosynthetic eukaryotes that are responsible for about 20% of photosynthesis on Earth [1]. Such phototrophic organisms are key players in shaping the global carbon cycle, as they can serve as food for higher eukaryotes and, more importantly in the context of the present work, for heterotrophic bacteria. This latter group of bacteria includes ubiquitous microorganisms that usually thrive on organic carbon compounds produced by diatoms (or other phototrophs), thus recycling them back to carbon dioxide and water. Indeed, from a metabolic perspective, the phycosphere (the region that extends outward from an algal cell [5]) is characterized by an intense activity. This includes commensalism based on organic carbon supplied by phytoplankton and competition for mineral nutrients [14]. In particular, DOM release by diatoms (a widespread process that may result in up to 50% of the fixed carbon leaving the cell) represents the main nutrient source for heterotrophic bacteria occupying the phycosphere [30, 31, 29]. The chemical nature and concentration of the released compounds vary with phytoplankton species and the physiological status of the phytoplankton [9, 8]. However, although phytoplankton releases a broad range of organic compounds, amino acids, carbohydrates and fatty acids arguably represent the main components of DOM [40]. On the other hand, diatoms and bacteria compete for the usage of several inorganic ions. Studies concerning the assimilation of inorganic phosphate, for example, have shown that bacteria can compete with phytoplankton and that this potential competition can be nutrient concentration dependent [11, 39, 14].

Phototroph-heterotroph interactions can be tackled both experimentally and theoretically. Co-cultures embedding these two types of microorganisms represent excellent platforms for identifying possible metabolic cross-talks and other interactions between these two organisms [42, 23]. Complementary, theoretical models have allowed shedding light on system-level characteristics of this biological association [37, 4, 11]. This latter approach has been mainly based on the use of systems of ordinary differential equations (ODEs) to predict the variation in time of phytoplankton and bacteria cell number in simulated co-culture experiments. In such systems, biological entities (e.g. bacterial and/or phytoplankton cells) are represented as discrete units, whose concentration in time (i.e. growth rate) is a function of a few parameters such as nutrients concentration and/or the growth rate of the eventual competitor.

The increasingly popular genome-scale metabolic modelling (GSMM) approach offers the opportunity to examine the phototroph-heterotroph association (and ecosystems in general [38, 10]) under a different perspective and with higher resolution. Indeed, despite being demanding, the assembly of genome-scale metabolic reconstructions accounting for the (almost) entire set of metabolic reactions of a cell is today feasible in a reasonable amount of time and for virtually every strain whose genome has been sequenced [22]. Additionally, the use of constraint-based metabolic modelling tools (e.g. flux balance analysis, FBA) allows simulating metabolic fluxes at the whole cell level and predict realistic growth rates, given a list of nutrients available to the cell and few biologically sound assumptions (e.g. a cellular objective function and reaction bounds). To date, however, this approach has never been exploited to systemically investigate the metabolic cross-talk between diatom and bacterial representatives. The aim of this work relies in the reconstruction of an integrated, two-organism metabolic model accounting for the metabolic cross-talk known to occur between phototrophic and heterotrophic microbes and in its exploitation for *in silico* reproducing the behaviour of this biological system. This was accomplished using available resources for two model organisms, i.e. the heterotrophic bacterium *Pseu-doalteromonas haloplanktins* TAC125 (*Ph*TAC125) and the phototrophic diatom *Phaeodactylum tricornutum* (*P*TRI). Our group has recently assembled and experimentally validated a genome-scale metabolic reconstruction for the heterotrophic Antarctic bacterium *P. haloplanktins* TAC125 [19]. This microorganism is considered the model of cold-adapted bacteria and a growing body of literature has illustrated its main physiological features [43, 35]. Similarly, *P. tricornutum* is considered one of the model diatom representatives and many experiments have contributed to the deep characterization of its physiology available nowadays. Genome-scale metabolic reconstructions are available for this diatom, the most recent dated back to the early 2017 [25]. Importantly, representatives of these taxa have been shown to co-occur in oceanic samples [28].

We here show that this combined metabolic reconstruction is able both to recapitulate the main known features of this simple microbial community (including competition and commensalism) and to dynamically represent/predict the behaviour of this biological system. Being based on genome-scale reconstructions, our approach allows exploring and interrogating the model at the single metabolic reaction/metabolite level. Further, using the fixed points predicted by CBMM, a simpler ODEs model was implemented and used to model the dynamics of the *Ph*TAC125-PTRI co-culture and to perform stochastic modelling of this synthetic ecosystem. We anticipate that the implemented modelling framework will represent a valuable resource for a more focused experimental design to study diatom-bacteria interaction.

## 2. Methods

### 2.1. The constraint-based model

The diatom-bacterium synthetic ecosystem was built starting from the single-organism models of *P. tricornutum* [24, 25] and *P. haloplanktis* TAC125 [19]. We adopted the following strategy to build a multi-compartment metabolic model accounting for the metabolic interactions of these two organisms:

1. As the two reconstructions encompassed different naming schemes, we mapped the two reconstructions to the same name space using the MetaNetX server [7].
2. To minimize the required interventions on the original reconstructions, only the names of the boundary metabolites common to the two reconstructions were changed to the corresponding MetaNetX code. In other words, only the set of metabolites that can be exported and imported by the two organisms were named according to the same name space. These (14) metabolites (Supplementary Material S1) will represent the pool of nutrients that can be eventually shared among the two organisms by means of different interaction types (namely, competition and cooperation).
3. A multi-compartment reconstruction was built. This included a compartment (s) in which the two single-organism reconstructions are inserted and in which extracellular metabolites can be secreted and/or up taken. This compartment ideally recapitulates the growth medium. Exchange reactions regulating the amount of nutrients present in this common compartment were included in the overall reconstruction to modulate their availability to the two organisms. Additional compartments included: extracellular and cytoplasmic bacterial compartments (z and t, respectively), extracellular, cytoplasmic, chloroplastic, mito-condrial, thylakoid lumen and peroxisomic diatom compartments (e, c, h, m, u, x, respectively). A schematic representation of the multicompartment model is reported in Figure 1.

**Figure 1:**
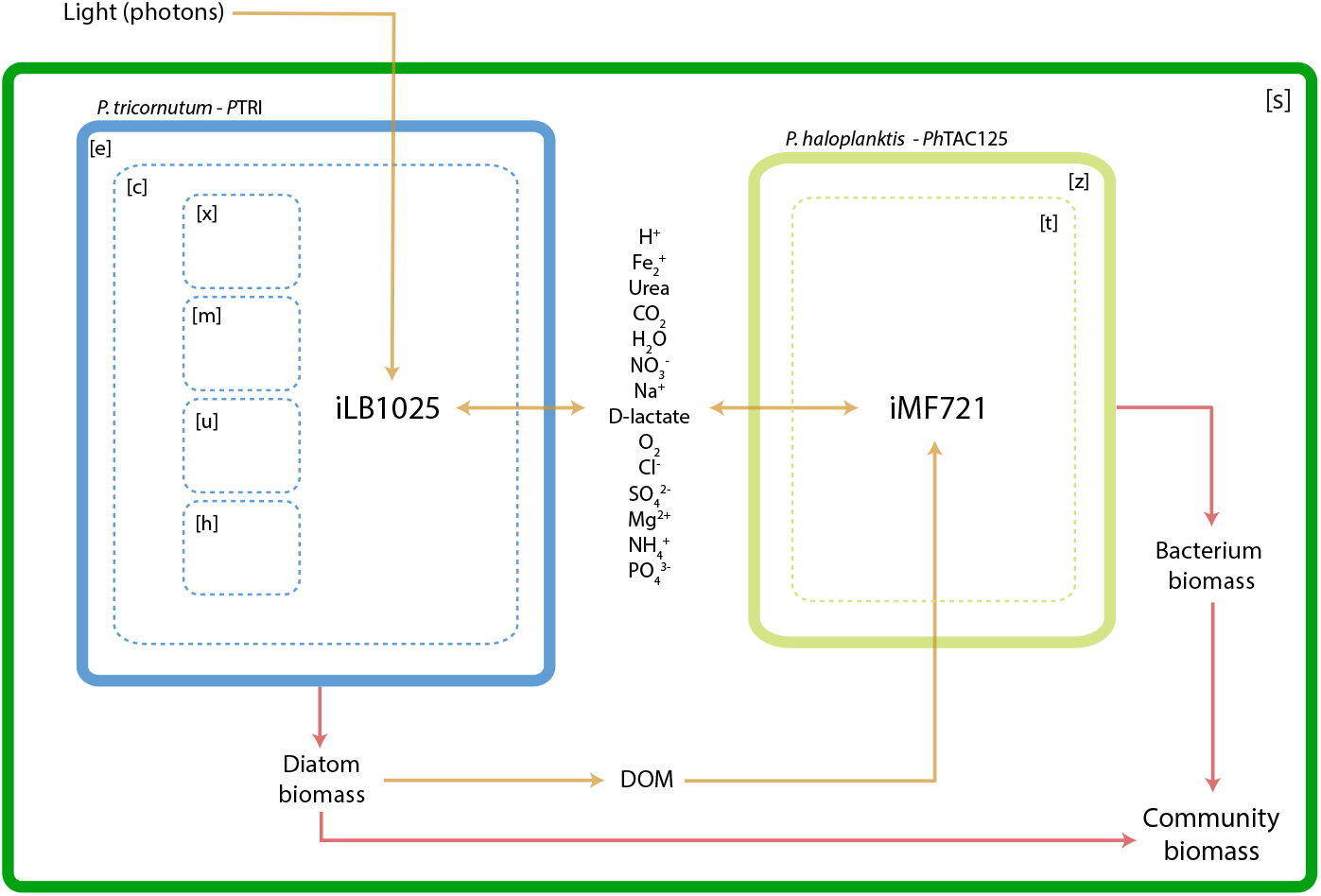
A) Schematic representation of the integrated diatom-bacterium metabolic reconstruction used for constraint-based metabolic modelling.

Concerning community biomass, we preliminary defined a community-level objective function by linearly merging the species’ biomass equations into the following biomass generating reaction:

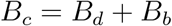

where *B_c_, B_d_* and *B_b_* represent community, diatom and bacterial biomasses, respectively.

In our integrated reconstruction, DOM is secreted by the diatom and represents the only C source for the bacterium. More in detail, we introduced a DOM release reaction in the combined model, accounting for the exudation of a fraction (10%) of the biomass produced by the diatom through auxotrophic growth. Following experimental evidences, the composition of the secreted DOM included carbohydrates (glucose, mannose, arabinose, xylose, fucose, galactose and glucuronate), fatty acids (myristic acid, palmitic acid, stearic acid, palmitoleic acid, alpha-linoleic acid, eicosapentanoic acid) and all the 20 amino acids; the kind and the relative amount of each single compound released in the external medium by the diatom was derived from the biomass composition equation of the original diatom metabolic reconstruction and/or from specific literature [25, 6]. See Supplementary Material S1 for the specific model reactions accounting for DOM release.

Concerning *Ph*TAC125 carbon uptake, it is well-established that, when grown in a complex medium, i) the metabolism of this bacterium is geared towards the consumption of amino acids [27, 18, 35, 43] and ii) the bacterium thrives on a very narrow spectrum of carbohydrates [19]. We then assumed that the bacterium would preferentially feed on a reduced set of DOM-derived compounds rather than on the entire set of single DOM constituents listed above. In particular, when facing a complex mixture of amino acids, *Ph*TAC125 selectively starts feeding on glutamate, serine, threonine, asparagine and aspartate; the remaining amino acids are consumed only after the exhaustion of this set [43]. We then assumed that, out of all the compounds released by the diatom, the bacterium would only feed on these amino acids.

Lower bounds of amino acids uptake reactions of *Ph*TAC125 were initially set to −1 *mmol/g * h*^−1^ as this was shown to resemble the growth phenotype of this bacterium when growing in an amino acids rich medium [19]. The other boundaries of the integrated model were set as described in Supplementary Material S1.

The integrated *Ph*TAC125-PTRI metabolic reconstruction was next converted into a dynamic, constraint-based, model of this co-culture using the approach implemented in DFBAlab [21] as described in detail in Supplementary Material S1. Mirroring the previously described constraint-based static model, *P*TRI thrives only on photons and *CO*_2_ and releases DOM in the culture; DOM, in turn, represents the only C source for *Ph*TAC125. Further, the growth of both organisms is dependent on phosphate and accordingly, they compete for it. All the other micro-nutrients were considered to be present in non-limiting concentrations (i.e. the lower bounds of the corresponding reactions were set to −1000). A negligible initial amount (1×10^−5^ *mmol/l*) of the other nutrients whose concentrations were allowed to change in time (e.g. DOM components) was assumed, to ensure their accumulation during the dynamic simulation (non-zero derivatives). A set of differential equations was then used to model the dynamic co-growth of *Ph*TAC125 and *P*TRI and the consumption of the nutrients in the medium (see Supplementary Material S1).

### 2.2. The stochastic model of the synthetic ecosystem

The dynamic description of the *Ph*TAC125-PTRI co-culture was also formulated as an ensemble of stochastic rules using a set of chemical equations. To this end, we lumped all the species of the constraint-based model and denote by *D, B, C* and *P* one individual of diatom, bacterium, carbon and phosphate, respectively. In this system (mimicking the previous set up) the diatom relies on phosphate and light to produce biomass and dissolved carbon that, in turn, represents the only carbon source for the bacterium. As we decided not to include (the constant inflow of) photons in this ODEs dynamic system of equations, the equation describing the growth of diatom in the system can be written as:

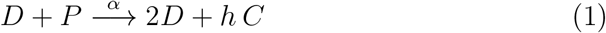

where *α* represents the overall rate of the reaction, while *h* is the number of produced molecules of carbon.

Bacteria in the system relies on dissolved carbon (produced exclusively by the diatom) and phosphate to duplicate and eventually increase their biomass. We describe this relationship using the following equation:

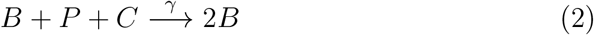

where *γ* represents the overall rate of the reaction.

Similar to [11], the growth rate of the diatom is a function of *P* and *D*, whereas bacterial growth depends on *P* and *C* availability.

Additionally, phosphate and carbon are injected into the system at a constant rate

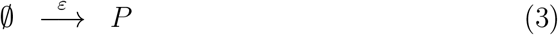

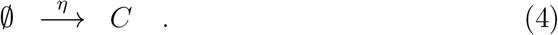

It is to be noted that, in our synthetic ecosystem, C can only be produced by the activity of the diatom and no influx of external C was allowed. For this reason the parameter regulating the inflow of C(*η*) was set to 0 during the dynamic simulations.

Eventually, all the species undergo degradation according to

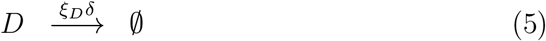

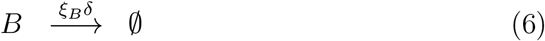

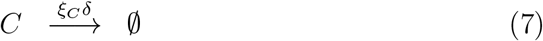

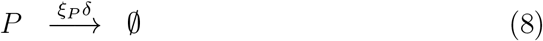

where *δ* is the degradation rate and *ξ_i_* with *i* = *D,B,C,P* are diluition rate constants.

At each time *t* the system is fully described by **n** = (*n_D_, n_B_, n_C_,n_P_*), a vector whose components *n_i_* for *i* = *D,B,C,P* are discrete quantities that identify the total number of elements of species *i*. To proceed in the analysis, it is necessary to introduce the transition rates *T*(**n**′|**n**) from the state **n** to the new state **n**′. These transitions originate from Eq. (1)–(8) and their explicit forms can be found in in Section 1.4 of Supplementary Information S1. In this way, the dynamics of the system is governed by a master equation whose generic form reads

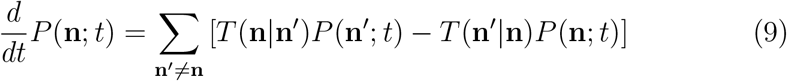

where *P*(**n**; *t*) labels the probability for the system to be in state **n** at time *t*. Even though in most situations this equation does not allow a closed-form solution, it represents a powerful tool that enables one to get insight into the traditional continuous and deterministic rate equations as well as fluctuations stemming from the discreteness of the system.

Let us first focus on the deterministic case. Following the detailed analytic derivation described in Section 1.4 of Supplementary Information S1, it can be proved that, in the limit where the volume of the system goes to infinity [41], the stochastic model described by Eq. (1)–(8) can be mapped into the following set of ordinary differential equations

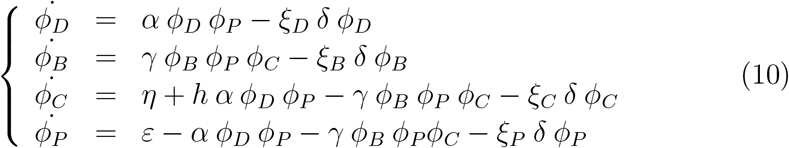

where *ϕ_D_, ϕ_B_, ϕ_C_* and *ϕ_P_* stand for the continuous concentrations associated to diatom, bacterium, carbon and phosphate, respectively. Setting equal to zero the temporal derivatives appearing on the left hand side of Eq. (10), the solution of the corresponding nonlinear system gives the equilibrium point that we here label 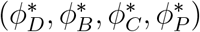. Its components are

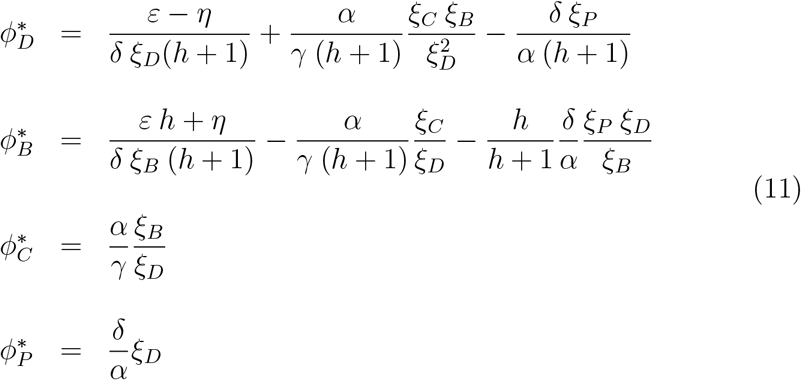

Since the ODEs model mimics the mechanism underling our dFBA implementation, we expect that the stationary concentrations resulting from the two different approaches to be the same. For this reason, denoting by *FBA_D_, FBA_B_, FBA_C_* and *FBA_P_* the stationary concentrations of, respectively, diatom, bacterium, carbon and phosphate resulting from the dFBA simulation, our idea is to set them equal to 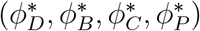. This procedure yields a non linear system that can be analytically solved to find an estimation of the parameters, provided *ε, η, ξ_D_, ξ_B_, ξ_C_* and *ξ_P_* are known. In formulae:

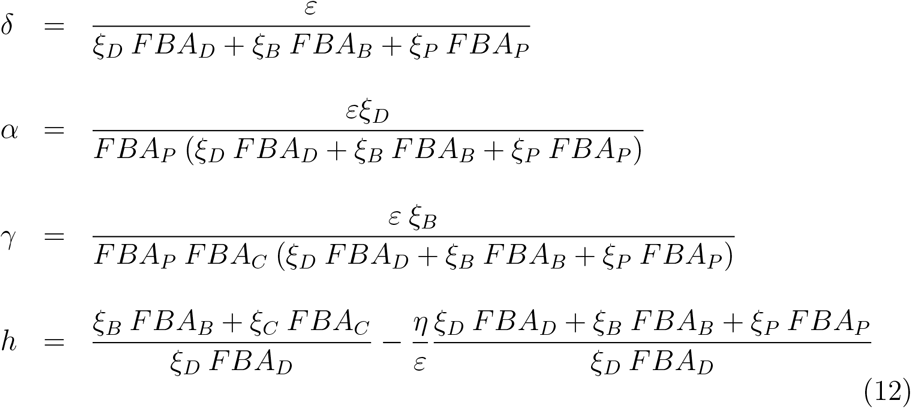

### 2.3. Modelling

Constraint-based simulations (FBA) were performed using COBRA toolbox v. 2.0.6 in MATLAB 2015b with Gurobi v. 6.5.0 as the linear programming solver. Dynamic FBA was performed using an *ad hoc* modified version of DFBAlab [21]. Stochastic simulations were performed using the Gillespie algorithm [20], essentially based on a dynamic Monte Carlo method. All the codes used in this work are available at https://synml.sourceforge.io

## 3. Results

### 3.1. The integrated reconstruction recapitulates phototroph-heterotroph metabolic cross-talk

In this work, we have combined two existing metabolic reconstructions (namely iLPB1025 and iMF721) in a single, integrated model to simulate a synthetic ecosystem characterized by (the metabolic) interactions existing between a phototroph (the model diatom *P. tricornutum* and a heterotrophic bacterium (*P. haloplanktis* TAC125). The integrated metabolic model consists of 1746 genes, 3588 reactions and 2861 metabolites. This synthetic ecosystem encompasses 9 compartments, including 1 external, shared compartment where cellular products are secreted/imported by the two organisms, 6 and 2 cellular compartments belonging to the diatom and the bacterium reconstructions, respectively. In an initial phase, reaction boundaries were defined as reported in Methods. A community biomass objective function was defined as the linear combination of *P*TRI and *Ph*TAC125 biomasses, as described in Methods. An FBA simulation was performed assuming such conditions, allowing a preliminary outlook on the metabolic phenotype of the diatom-bacterium community. This analysis revealed that the model correctly predicts the main known features for such a biological system in that the diatom is predicted to uptake *CO*_2_ from the environment and to use 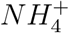 as *N* source. Consistently, *O*_2_ is released in the environment as a product of the diatom metabolism. The simulated conditions support the growth (and *O*_2_ consumption) of the heterotrophic bacterium *Ph*TAC125, whose predicted growth rate is 0.02 *h*^−1^. No ammonia uptake is predicted for the bacterium, consistently with the fact that amino acids (part of the DOM fraction) can be used as both C and N source by *Ph*TAC125. We then checked the metabolic interdependencies existing between the two models in the ecosystem. To this aim, we computed the Pareto front between diatom and bacterium growth rates (Figure 2a). In its simplest formulation, the Pareto front represents the trade-off between the two organisms maximal growth rates. Accordingly, the surface delimited by the Pareto optimality frontier identifies the set of optimal solution of our double optimization (diatom and bacterium growth). Such solutions (i.e. a couple of growth rate values) are said to be Pareto-optimal as there are no other solutions that can better satisfy all of the objectives simultaneously. In particular, Figure 2a revealed three main features: i) a situation in which both the diatom and the bacterium growth at their corresponding highest achievable growth rate is not included in the plane of feasible solutions. ii) Diatom growth is predicted even in the absence of bacterial growth but, conversely, iii) bacterial growth in the absence of diatom growth is not a feasible solution. Overall, this analysis confirmed that the two models embedded in the integrated reconstruction are not independent of each other and that both competition (organisms’ growth rates are independent one another and infeasible simultaneous bacterial and diatom maximal growth) and commensalism (no bacterial growth in the absence of diatom growth) can be described by our system. The latter feature is clearly explained by the fact that diatom-derived DOM is the only carbon source that is made available to the bacterium. Indeed, decreasing the amount of light available to the diatom resulted in a decrease of the overall community growth rate (Figure 2b). No clues concerning which substrate(s) is responsible for the competition between the two organisms can be derived simply by inspecting the Pareto frontier. To identify those shared metabolites whose availability limited the growth of both organisms, we performed a robustness analysis, i.e. we evaluated the effect on *Ph*TAC125 and *P*TRI growth rates of perturbing the uptake rate of each of the (14) compounds that can be taken up by each organism in the reconstruction. We identified two macronutrients whose uptake is essential for sustaining the growth of both organisms, namely phosphate and sulphate. As shown in Figure 2c and d, perturbing the uptake rates of such compounds strongly influences the growth rate of *P*TRI and *Ph*TAC125 and no growth is predicted when their uptake rate is set to 0. As outlined below, metabolic competition over these inorganic ions is a well-described feature among representatives of diatoms and bacteria [16, 11].

**Figure 2:**
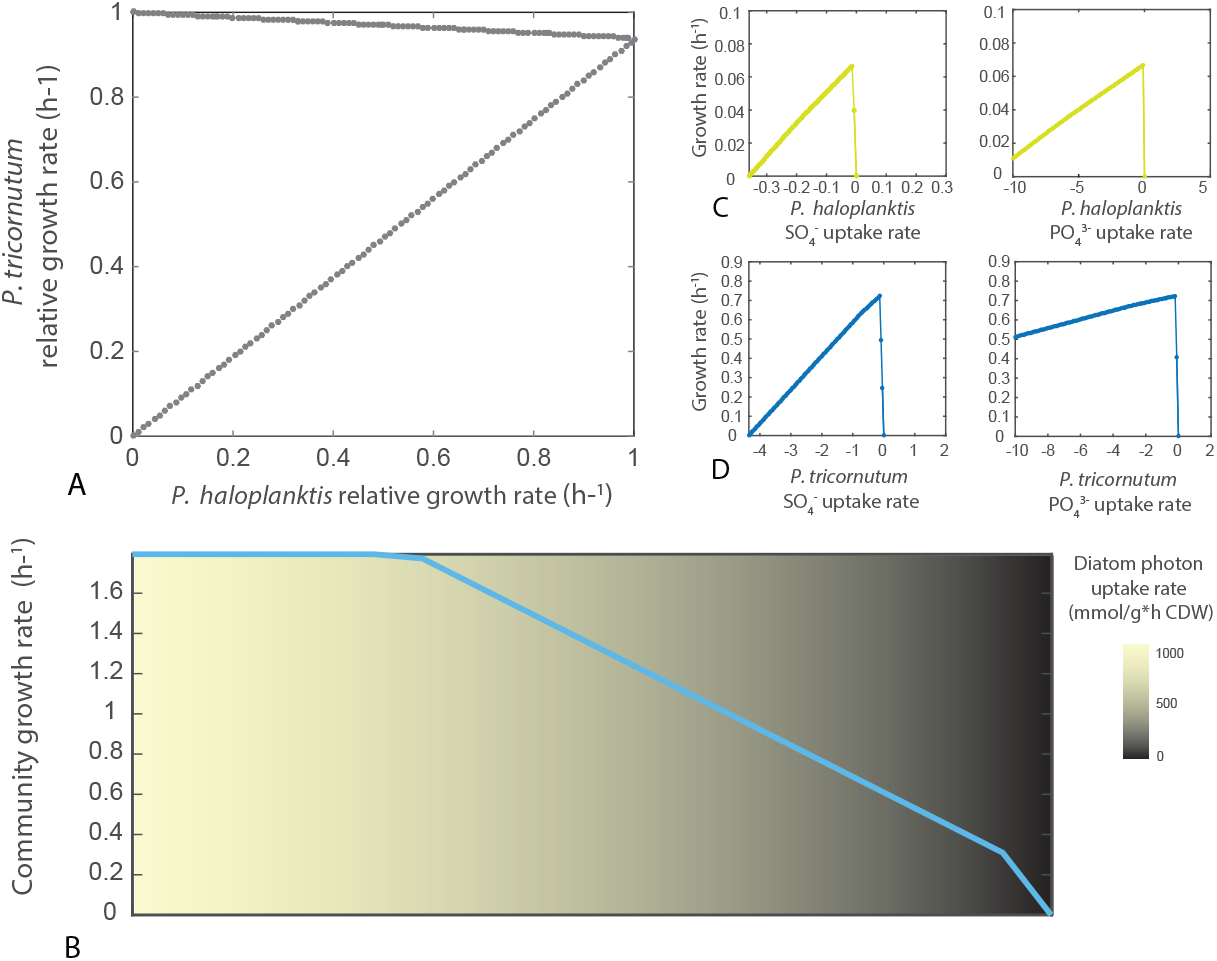
a) Pareto front between microbial and diatom growth. Growth rates are expressed relatively to maximum growth achievable by each microorganism. b) Community biomass at decreasing light availability. c) Robustness analysis showing the relationship between bacterium growth rate and different simulated phosphate uptake-fluxes. d) Robustness analysis showing the relationship between diatom and bacterium growth in relation to different simulated sulphate uptake-fluxes.

### 3.2. A dynamic FBA model for the phototroph-heterotroph metabolic cross-talk

To understand the dynamics of the metabolic interactions between *Ph* TAC125 and PTRI, we implemented a dynamic, constraint-based model of their growth in a well-stirred reactor as described in [21]. Mirroring the simulation previously described, *P*TRI thrives only on photons and *CO*_2_ and releases DOM in the culture; DOM, in turn, represents the only C source for *Ph*TAC125. The amount of matter released by diatom in the form of DOM is highly variable and seems to be dependent on the species and the environment considered (see below); in this initial simulation we assumed that 10% of the fixed C was released in the culture and could be metabolised by the bacterium. We started the dynamic simulation from a 1:1 ratio in *Ph*TAC125 and *P*TRI cell concentrations in this synthetic community (0.01 *gCDW/L*). We then set an initial, non limiting, phosphate and C concentration of 5 *mmol/L* and 10 *mmol/L*, respectively; additionally, the concentration of the inlet flow of phosphate (*δ*) was set to 0.0016 *mmol/L*, mimicking the observed concentration of phosphate in natural conditions [45]. As *Ph*TAC125 metabolism is known to be geared towards the consumption of amino acids [27, 18, 35, 43], we monitored the concentration of the exchanged amino acids (glutamate, serine, threonine, asparagine and aspartate) across the entire simulation.

As shown in Figure 3b and c, both organisms start growing thanks to the photons income (the diatom) and to the initial amount of C present in the system (the bacterium) (Figure 3c and d). The ecosystem growth, however, is interrupted by the exhaustion of the phosphate excess in the medium (Figure 3a). The diatom starts a second growth phase growing earlier than the bacterium, compatible with the fact that the latter has no carbon source available until *P*TRI has reached a certain cell density and has released a sufficient amount of DOM-derived nutrients in the medium. After the diatom production of DOM is sufficiently high (Figure 3d), the bacterium starts another growth phase (Figure 3c) which is further interrupted by phosphate decrease. Afterwards, as phosphate is added to the co-culture in a limiting amount, an oscillatory pattern is observed, reflecting the competition for phosphate on one side and the fact the *Ph*TAC125 thrives only on released DOM on the other. Indeed, as DOM derived amino acids get more abundant, *Ph*TAC125 increases its phosphate requirement to sustain growth and consequently will reduce the amount of available phosphate for *P*TRI that, in turn, will slow down it’s growth rate and the amount of available C to the bacterium. Such an oscillatory pattern is clearly reflected when computing the ratio between bacterium and diatom biomass throughout the simulation (Figure 3a). The simulated culture conditions will then lead to a steady state in which the concentrations of the different species (both organisms and dissolved nutrients) won’t change over time. In our simulation, the equilibrium is reached with a *P*TRI concentration that is 100-fold higher than that of *Ph*TAC125. Additionally, the final diatom cellular concentration and the (average) predicted growth rate predicted by our simulation throughout the simulation were 0.038 *gDW/L* and 0.0128 *h*^−1^ for PTRI and 0.0025 *gDW/L* and 0.0164 *h*^−1^ for *Ph*TAC125.

**Figure 3:**
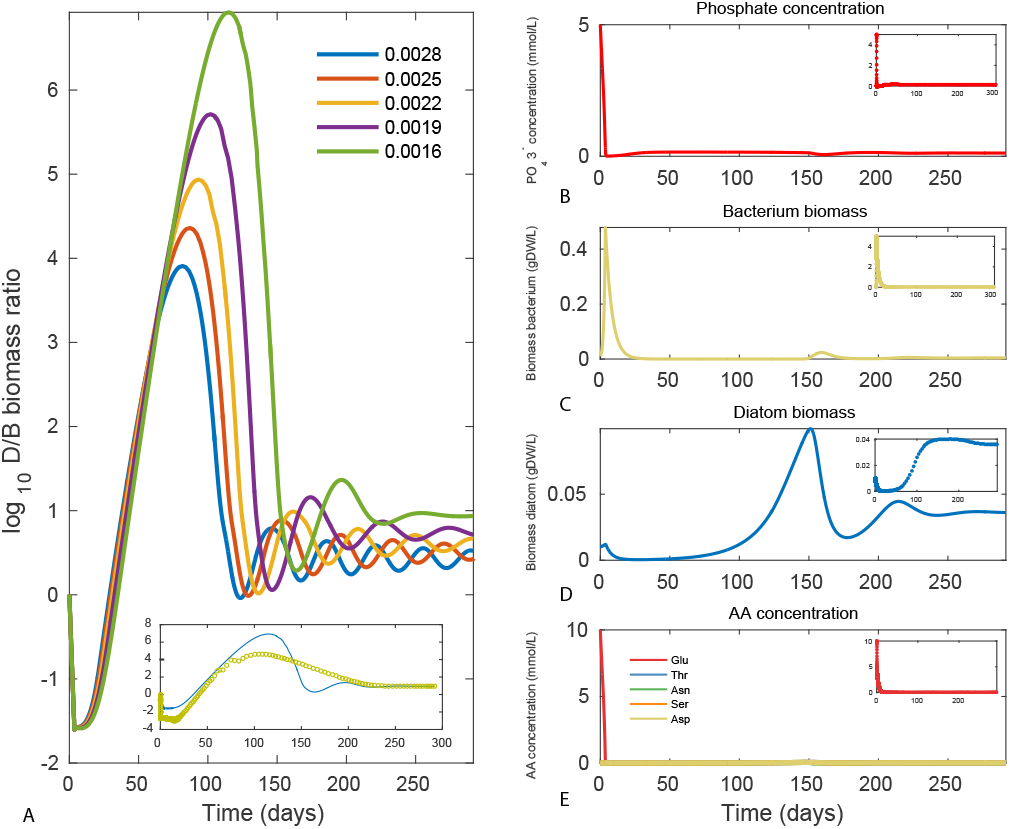
Dynamic simulation of the phototroph-heterotroph co-culture showing the change in time of the log_10_ of the diatom-bacterium biomass ratio (a), phosphate concentration (b), bacterial biomass (c), diatom biomass (d), amino acids concentration (e)

Increasing the concentration of phosphate available to the two organisms resulted in a longer and larger oscillatory pattern before the final steady state is reached (Figure 3a). Apparently, an increased phosphate concentration leads to a positive feedback according to which bacterial growth rate increases, DOM nutrients are consumed more rapidly and a stronger bacterium-diatom competition for phosphate is established. This trend is further amplified setting the dilution rate of phosphate equal to zero, allowing both organisms to consume all the phosphate that is actually introduced into the system. In this case all the species in the ecosystem display sustained oscillations (see Supplementary Information S1).

Clearly, the amount of DOM released by the diatom and used by the bacterium as carbon and energy sources is crucial for maintaining the equilibrium between *Ph*TAC125 and *P*TRI biomass concentration in this closed system. We monitored the effect of varying the amount of DOM released by the diatom on the stability of the co-culture system. Thus, we performed distinct dynamic simulations, varying for each of them the ratio between biomass accumulation and released DOM. Our analysis shows that, as it might expected, the *P*TRI/*Ph*TAC125 ratio in this synthetic ecosystem increases with the decrease of the released DOM; this reflects both the fact that, in these simulated conditions, the diatom redirects a greater fraction of cellular intermediates to the synthesis of biomass rather than to the secretion of C compounds and that, consequently, the bacterium has a lower amount of available nutrients (Figure 4).

**Figure 4:**
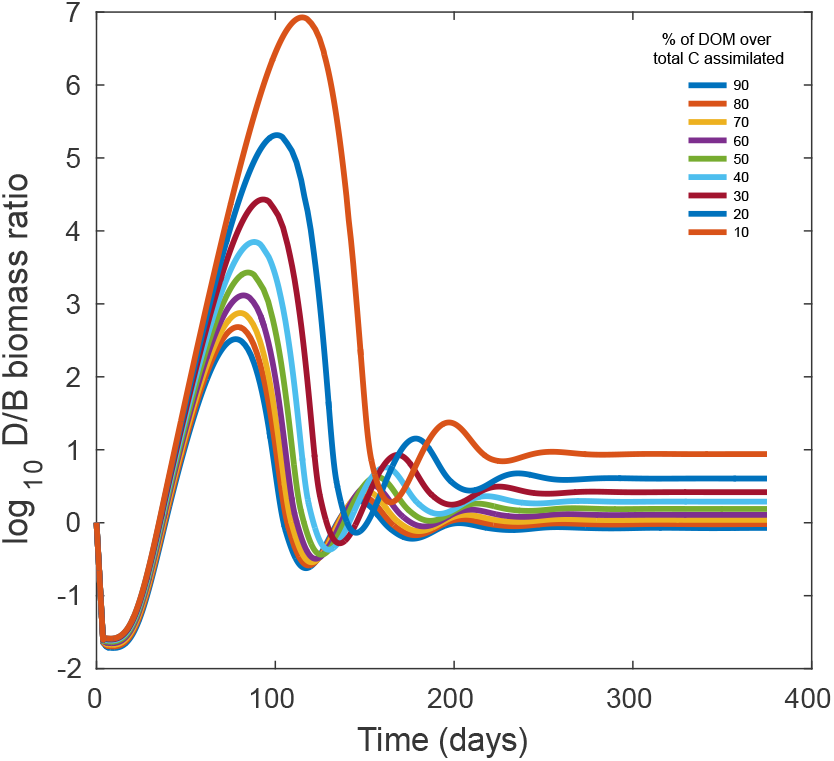
Dynamic simulation of the diatom/bacterium (D/B) biomass ratio at different fractions of diatom-released DOM expressed as percentage of the total fixed carbon

### 3.3. ODE-based modelling of iDBE

As shown, the dFBA model recapitulates the behaviour of a *Ph*TAC125-*P*TRI co-culture. Nevertheless, this dynamic modelling framework is purely deterministic and, as such, it cannot deal with possible noise and fluctuations of the surrounding environment. Indeed, it is widely accepted that marine environment is subjected to recurrent fluctuations in nutrient levels through the movement of tides and currents [13]. For this reason, the next step was to use the results of this dynamic simulation to set up a simpler and more controllable dynamic model based on ODEs (see Methods). Using the fixed-point values obtained for each parameter, we performed a dynamic simulation of the diatom-bacterium co-culture using Eq. (10) to check whether, despite being computationally simpler, the implemented ODEs model could qualitatively reproduce the behaviour of the dynamic constraint-based modelling. As shown in the insets of Figure 3a-e the ODEs-based model correctly predicts an initial increase of bacterial, diatom and carbon species; also, in agreement with the previous simulation, phosphate is predicted to decrease from its initial concentration determining the stop of the growth of the aforementioned species. Similarly, the biomass ratio of diatom and bacterial cells predicted by the ODEs-based model qualitatively resembles that resulting from the constraint-based model, displaying an initial predominance of bacterial cells followed a rapid and persistent shift towards a diatom-dominated culture. As expected, the ODEs-based model then approaches the same steady point predicted by the dFBA model, although displaying a reduced oscillatory trend. This is likely due to the difference in the basic formulation of the two models, with the dFBA one embedding more variables and parameters and thus capable of accounting for more complex behaviours of this specific co-culture.

After checking that the simpler dynamic system displayed the same over-all trend of the dFBA model, we wanted to address the possible effects of fluctuations on the simulated metabolic cross-talk between *P*TRI and *Ph*TAC125. The mathematical models described previously have approached the problem via deterministic paradigms, overlooking the possible role played by the noise, intrinsic to the phenomenon under investigation. These aspects become particularly important when, as in this case, accounting for the presence of diverse chemical species in a spatially diffusive environment. For example, the two organisms may compete with the nutrients (e.g. phosphate) and, at the microscopic scale, they might be hindering each other the binding and the uptake of the nutrient molecule. Under specific conditions (i.e. nutrients paucity and/or unbalanced biomass ration in the culture), such competition sustained by the stochastic component of the dynamics might result in different and alternative steady states.

In order to perform stochastic simulations, we initially adjusted some of the parameters of the model in order to get 1) a balanced ratio between diatom and bacterial cells at the fixed point and 2) concentration values for all the species embedded in the system sufficiently high to avoid their eventual extinction from the system after a few iterations. This involved reducing the dilution rate of *Ph*TAC125 and *P*TRI and increasing the fraction of DOM (in respect to produced biomass) leaving the diatom cell and representing the bacterial only C source (from 10% to 20%, Table 1).

**Table 1:** The main parameters used during deterministic and stochastic simulations

|  | Deterministic | Stochastic |
| --- | --- | --- |
| *PhTAC125* initial biomass (gDW/L) | .1 | 1 |
| *PTRI* initial biomass (gDW/L) | .1 | 1 |
| Phosphate initial concentration (mmol/L) | 5 | 5 |
| Carbon initial concentration (mmol/L) | 10 | 10 |
| DOM fraction | 10% | 20% |
| Dilution rate *PhTAC125* | 100% | 50% |
| Dilution rate *PTRI* | 100% | 10% |

As already stated, the master equation 9 cannot be solved and thus analytical progress is possible only making use of expansion techniques that requires specific assumptions. One available tool to overcome this limitation is to solve the stochastic model through dedicated numerical schemes such as the Gillespie algorithm [20]. A trajectory corresponding to a single Gille-spie simulation exactly belongs to the probability distribution solution of the master equation.

Assuming the initial conditions listed in Table 1, our analysis showed a little effect of noise on the stability of the synthetic ecosystem, with the two species displaying the same overall dynamics of the deterministic model (Figure 5). We also performed the same stochastic simulation for different values of the inlet phosphate concentrations *ϵ*, revealing no major influence of such parameter on the overall stability of the system (Supplementary File S1).

**Figure 5:**
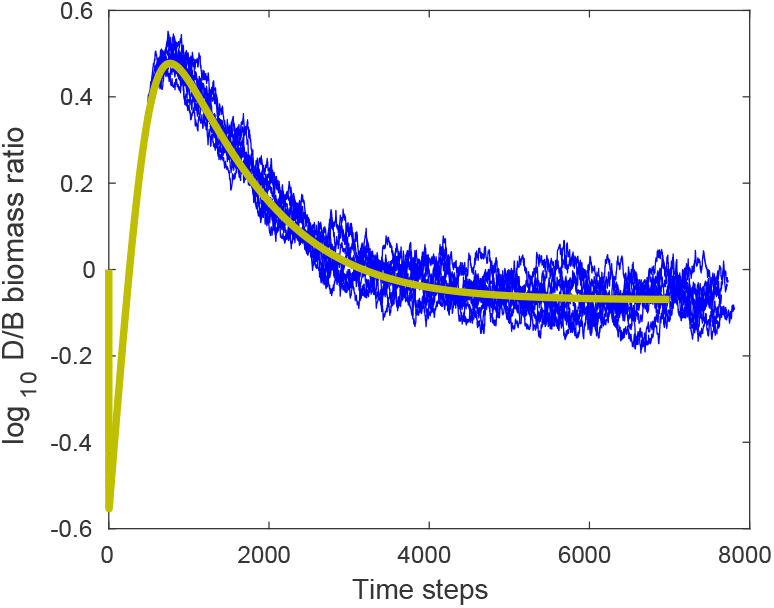
Stochastic simulation of the diatom-bacterium co-culture with fixed phosphate inlet and dilution rate. Each blue line represents a single simulation. The yellow line represents the dynamic deterministic simulation.

The other parameter that might influence the dynamics of the the synthetic ecosystem is the dilution rate of the diatom (*ξ_D_*) and the bacterium (*ξ_B_*) in the simulated bioreactor as it ultimately regulates the number of cells populating the co-culture. Thus, we tested the robustness of the system in respect to stochastic fluctuations for a different range of *FBA_D_/FBA_B_* cell concentration ratios. We performed a new series of stochastic simulations setting the dilution rates to 10, 20, 40, 60 and 80% for for both the diatom and the bacterium and tested all the possible combinations thereof. When dilution rates for both organisms are above 50% and the ratio between diatom and bacteria dilution rates is high (i.e. the diatom cell concentration does not sensibly exceed the bacterial one), the system displays a rather robust trend in respect to noise. Indeed, no major discrepancies compared to the deterministic simulation are observed (Figure 6a). At higher values of bacterial dilution rates, i.e. with lower bacterial cells in the co-culture, the equilibrium becomes rather unstable. When the ratio between diatom and bacteria dilution rates approaches 0.3 (i.e. bacteria die roughly three times faster than diatoms) some stochastic simulations deviate from the fixed point and, more specifically, the culture becomes diatom-dominated (Figure 6b). A further increase in bacterial dilution rate leads to a clear prevalence of diatom-dominated co-cultures among the whole set of stochastic simulations (Figure 6c) as virtually all the simulations lead to the maximum biomass concentration achievable by the diatom. As expected, we observed no bacterial-dominated co-cultures among all the stochastic simulations performed as the diatom-produced DOM is essential for *Ph*TAC125 survival.

**Figure 6:**
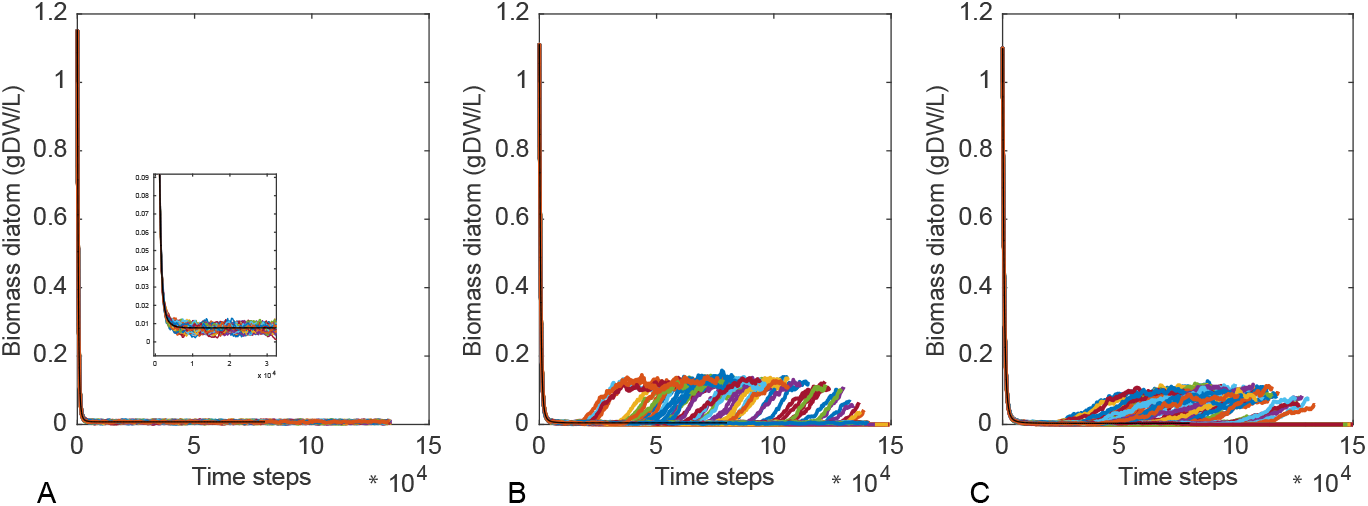
Diatom cell concentration at different dilution rates for diatom and bacterial cells. A) dilution rate ratio set to 0.5, B) 0.16 and C) 0.125). Each line represents a single stochastic simulation of the co-culture behaviour.

## 4. Discussion

In this work we have used two established genome-scale metabolic reconstructions to build a synthetic ecosystem accounting for (part of) the metabolic interdependencies between diatoms and bacteria. Applying CBMM (namely FBA) revealed that our model is able to capture the basic metabolic relationships as competition and commensalism, which are commonly observed in diatom-heterotroph associations [1]. In our synthetic ecosystem, the growth of the bacterium is totally dependent on the release of DOM by the diatom; on the other hand, the growth of the bacterium reduces the availability of inorganic ions (phosphate and sulfate) to the diatom, thus negatively affecting its growth.

The integrated reconstruction was then analysed by means of a dynamic framework, that is dFBA. This allowed the simulation of a co-culture system (stirred reactor) and to evaluate the effect of specific parameters (e.g. phosphate inflow concentration) on the dynamics of this biological system. In agreement with previous results from experimental diatom-bacteria cocultures and mathematical models, our simulations reached an equilibrium with the diatoms and the bacteria co-existing. By adjusting the different parameters encompassed by our model we showed that this equilibrium can be shifted towards bacterial or algal dominance and/or leading to interesting collective phenotypes (e.g. sustained oscillations). The final cellular concentration of the two organisms fall in the range of measured values for *P*TRI; indeed, a mono-culture of *P*TRI on a f/2 medium reached a final cell concentration between 150 and 200 *mg* and a *μ* of 0.025 *h*^−1^ (an average *μ* of 0.0128 *h*^−1^ was predicted by our simulation throughout the simulation) [44]. This relationship between *Ph*TAC125 and *P*TRI biomass concentration resembles the one observed when growing other representatives of these two groups of organisms in a laboratory setting [14]. Varying the concentration of the phosphate inflow (Figure 3) had an effect on the intermediate stage of growth with smaller but longer oscillations observed when the phosphate concentration is increased. Such collective oscillations are typically observed in the presence of metabolic co-dependencies [26]. We then tested the effect of varying the proportion of fixed carbon that is redirected by the diatom towards DOM secretion rather than biomass assembly on the overall balance of the entire system. Phytoplankton exudation and/or cell lysis represents a primary resource for bacterial metabolism [3]. The quantification of dissolved organic substances released by phytoplankton has long been the subject of research and debate; a few works based on extensive data re-analysis identified an approximative average range of 10 to 20% released DOM over a diatom total primary production, with values up to 50% and as low as 4% [40, 32]. Our model predicts a shift towards a diatom-dominated culture parallel to the decrease of the fraction of C secreted in the medium rather than redirected towards diatom biomass assembly (Figure 4). However, the two species coexist with all the DOM release rates tested comprised this interval.

Finally, using the fixed points predicted by the dFBA simulation, we implemented a stochastic dynamic model of the same *Ph*TAC125-PTRI coculture. This system is composed by 4 ordinary differential equations (Eq. (10)), accounting for the change in time of the concentration of *P*TRI and *Ph*TAC125 cells, DOM and phosphate. After successfully checking that the dynamic model resembled the trends observed during dFBA simulations, we tested the effect of noise on the equilibrium between the two microorganisms. We found the synthetic ecosystem to be quite robust at similar diatom/bacterial cell concentrations and for different phosphate inflows. In this case, every stochastic simulation resembled the trend obtained with the deterministic model. However, decreasing the ratio of diatom/bacterial cell concentration down to 0.3 (by regulating their corresponding dilution rates) resulted in a bistable ecosystem phenotype, with a few stochastic simulations leading to an entirely diatom-dominated co-culture. Interestingly, bacteria have been shown to play a role in the explosion of algal population (bloom) in mesocosm experiments, e.g. by inhibiting diatom aggregation and prolonging the bloom [36, 34]. On the basis of the results of our model, we hypothesize that bacterial cell concentration (and its effect on the metabolic interactions with the diatom) may also have a role in determining the shift towards a diatom-dominated culture. Below a certain threshold, stochastic noise may induce the extinction of bacteria from the co-culture and, as a consequence, reduce the overall competition with diatom for essential nutrients (e.g. phosphate). This, in turn, may lead to the explosion of the diatom population. This is in agreement with previous observations on the abrupt decrease of bacterial abundance concomitantly with algal blooms [34] and with the bistable behaviour of modelled planktonic systems [15].

## 5. Conclusions

Here we have presented a synthetic bacterial-diatom ecosystem. The model was specifically designed for describing possible metabolic interactions occurring between an heterotrophic bacterium (*P. haloplanktis*) and a diatom (*P. tricornutum*). Such interactions were studied at different levels, using different modelling techniques, ranging from (static) constraint-based to (dynamic) stochastic modelling.

The model presented here provides a first attempt to formalize the possible metabolic cross-talk occurring in (a part of) the microbial loop. In the building up of our model we relied on several approximations. First, grazers (e.g. metazoans capable of feeding on microalgae and thus possibly contributing to their extinction from the synthetic ecosystem) were not accounted for by our metabolic reconstruction. To date, no genome-scale metabolic reconstruction exists for an organism that might resemble the features of a grazer inside the microbial loop; on the other hand, tuning the diatom dilution rates in the ODEs-based model may somehow account for an increased or decreased grazers’ activity in the ecosystem. Furthermore, in our proof-of-concept reconstruction we have identified possible metabolic interactions only on the basis of the genome sequence of the two microbial representatives. It is entirely possible that other metabolic cross-talks between these organisms exist and that the network of shared metabolites is more complex than the one hypothesized in this work. Metabolomic experiments conducted over co-cultures of *Ph*TAC125 and *P*TRI and their corresponding single cultures will clarify this point and highlight untapped metabolites exchange between these two organisms. Finally, at present, there are no direct evidences of the co-occurrencw of *P*TRI and *Ph*TAC125 in natual environment, despite *Pseudoalteromonadaceae* representatives have been shown to be the prevalent microbial family throughout most of the growth phases of the diatom [28]. The framework presented is currently driving the implementation of an experimental diatom-bacterium co-culture that, in turn, will be used to train our model and adjust all the parameters that are currently unknown. Once properly trained, it will be possible to reliably use our synthetic ecosystem to predict interspecies dynamics under numerous conditions. This might include, for example, the study of the effect of environmental perturbations on the overall system (e.g. climate change-related perturbations) or the possible exploitation of this ecosystem for biotechnological purposes.

